# The *Drosophila* Secretome

**DOI:** 10.64898/2026.08.24.746765

**Authors:** Yanhui Hu, Nahid Ausrafuggaman, Jenna Kalina, Austin Veal, Mujeeb Qadiri, Ah-Ram Kim, Norbert Perrimon

## Abstract

Secreted proteins are synthesized within the cell and actively transported outside, where they contribute to development, physiology, immunity, and cell–cell communication. In *Drosophila melanogaster*, many secreted proteins are evolutionarily conserved, making this species a powerful model for studying human development, cancer, neurobiology, and immune regulation. To assemble the *Drosophila* secretome, we systematically analyzed all fly protein sequences with computational tools, integrating FlyBase, UniProt and Gene Ontology annotations, and incorporating large-scale proteomics datasets. We identified 5,280 genes encoding putative secreted proteins, assigned confidence ranks and scores based on the type and strength of supporting evidence, and generated an online resource (www.flyrnai.org/apps/fly_secretome/) for exploring these data. Comparison with the human secretome shows that 56% of *Drosophila* secretome genes are conserved in the human genome and of these, 81% are annotated as secreted in human. In addition, comparison with *Drosophila* single-cell transcriptomic data revealed that secreted proteins are more tissue-specific than other genes. Finally, we demonstrate the utility of this resource by analyzing changes in expression of genes encoding putative secreted proteins during aging based on snRNA-seq datasets from the Aging Fly Cell Atlas.

## INTRODUCTION

Secreted proteins are produced by cells and then actively exported outside the cell. These proteins are typically synthesized in the endoplasmic reticulum, processed through the Golgi apparatus, and then released into the extracellular space through vesicular transport (Palade 1975). Secreted proteins in *Drosophila melanogaster* perform diverse and essential functions throughout development, adult metabolic homeostasis, immunity, and cell-cell communication (Vierstraete *et al*. 2005; Handke *et al*. 2013; Bosch *et al*. 2026; Yoshida *et al*. 2026). Because many signaling pathways are evolutionarily conserved, studies in *Drosophila melanogaster* have provided major insights into human development, cancer biology, neurobiology, and immune regulation (Pandey and Nichols 2011).

The secretome refers to the complete set of proteins secreted by a cell, tissue, or organism into the extracellular environment through classical or non-classical secretion pathways. Large-scale experimental approaches have identified hundreds of secreted proteins in *Drosophila* (Bosch *et al*. 2026) but a comprehensive analysis aimed at identifying all potential putative candidates is still missing. In this study, genes encoding putative secreted proteins, referred to as “secreted genes” for simplicity, annotated by FlyBase (Ozturk-Colak *et al*. 2024), UniProt (UniProt 2023) and Gene Ontology consortiums (Gene Ontology 2026) were integrated with large-scale proteomic datasets and all protein sequences of *Drosophila melanogaster* were analyzed with computation tools such as SignalP (Teufel *et al*. 2022) and DeepLoc (Nielsen 2025) algorithms to predict the presence of secretion signal peptides and subcellular localization, respectively. In total, 5,280 genes were identified as encoding putative secreted protein(s). Confidence was annotated based on the source as well as the number of sources providing experimental evidence. Further, we built an online resource (www.flyrnai.org/apps/fly_secretome/) to facilitate mining of the *Drosophila* secretome as well as relevant annotation and datasets.

Several bioinformatics analyses, including conservation analysis, were performed on the *Drosophila* secretome by comparing it to the secretomes assembled for other species, including human, mouse, rat, and zebrafish. Additionally, the transcript-level expression patterns of the *Drosophila* secretome in adult flies were analyzed using the Fly Cell Atlas (FCA) scRNA-seq dataset (Li *et al*. 2022). We demonstrate the utility of this resource by analyzing scRNA-seq transcriptomic datasets from Aging Fly Cell Atlas (AFCA) data (Lu *et al*. 2023) and surveyed the transcriptomic changes of secreted genes in various tissues during the aging process.

In summary, this study demonstrates an effort to build a comprehensive bioinformatics resource of *Drosophila* putative secretome to facilitate functional discovery in *Drosophila*.

## METHODS

### Assembly of the *Drosophila* Secretome

Gene group annotation of “receptor ligands” was retrieved from FlyBase (FB2026_03) (https://flybase.org/reports/FBgg0001105). At UniProt (https://www.uniprot.org/), the Advanced search option with the taxonomy filter [7227] and the subcellular location term “Secreted” [SL-0243] filter was used to obtain a set of UniProt accession numbers which were then mapped to FlyBase

GeneIDs using the ID mapping page at UniProt (https://www.uniprot.org/id-mapping). Genes annotated the gene ontology term “extracellular region” were retrieved from FlyBase using the Vocabularies search page (https://flybase.org/vocabularies). Secreted genes identified in a large-scale proteomics study were retrieved from a supplementary table 3B associated with the publication (Bosch *et al*. 2026). FlyBase GeneIDs from various sources were synchronized to FlyBase version FB2026_03 using FlyBase ID Validator tool (https://flybase.org/convert/id) and then integrated. The genes from these resources were designated the high rank tier of the *Drosophila* Secretome (Figure 1A).

**Figure 1.**
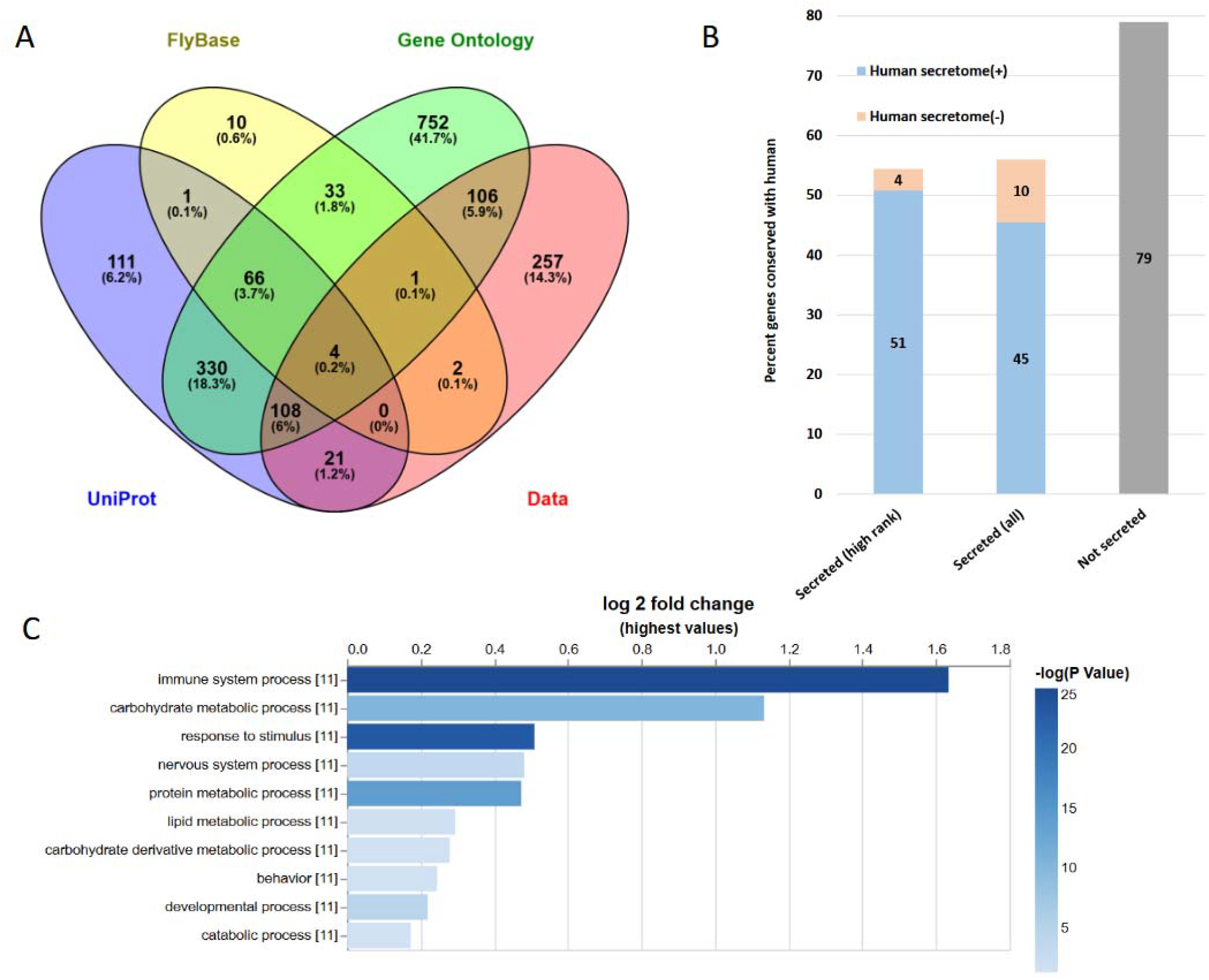
Assembly of a *Drosophila* Secretome. A.) Schema of gene selection from annotated resources. B.) Conservation of *Drosophila* secretome. C.) Gene set enrichment analysis of high rank secreted genes using PANGEA. The generic gene ontology slim terms of biological process were selected. The height of the bar graph reflects the log2 fold enrichment while the darkness of the color reflects P values (-log10 P value).

The FASTA file of the full *Drosophila* proteome was obtained from FlyBase ftp site (https://s3ftp.flybase.org/genomes/dmel/dmel_r6.69_FB2026_03/fasta/dmel-all-translation-r6.69.fasta.gz), which was used as the input file for various prediction algorithms. Software codes from SignalP 6.0 (Teufel *et al*. 2022) (https://services.healthtech.dtu.dk/services/SignalP-6.0/), TargetP 2.0 (Emanuelsson *et al*. 2000; Emanuelsson *et al*. 2007) (https://services.healthtech.dtu.dk/services/TargetP-2.0/), DeepLoc 2.0 (Nielsen 2025) (https://services.healthtech.dtu.dk/services/DeepLoc-2.0/), DeepTMHMM (Hallgren *et al*. 2022) (https://services.healthtech.dtu.dk/services/DeepTMHMM-1.0/), TMHMM 2.0 (Krogh *et al*. 2001) (https://services.healthtech.dtu.dk/services/TMHMM-2.0/), and ProP 1.0 (Duckert *et al*. 2004) (https://services.healthtech.dtu.dk/services/ProP-1.0/) were downloaded and run locally. Both fast and slow mode of SignalP 6.0 were run and the outputs were combined. Additional predictions were done by Phobius (Kall *et al*. 2007) (https://phobius.sbc.su.se/) and TOPCONS (Tsirigos *et al*. 2015) (https://topcons.cbr.su.se/) at the corresponding web portals. The prediction results for signal peptides and transmembrane (TM) helix were integrated respectively, and then the voting counts were calculated as well as integrated into the *Drosophila* Secretome database. Because four algorithms were used to predict the presence of signal peptide (SignalP, TargetP, Phobius and TOPCONS), the voting score for signal peptide ranges from 0 to 4; use of four TM-prediction tools (deepTMHMM, TMHMM2, Phobius and TOPCONS) yields a voting score ranging from 0 to 4. ProP predicts propeptide cleavage sites in eukaryotic protein sequences and DeepLoc 2.0 predicts eukaryotic protein subcellular localization using deep learning; results from these tools were also integrated into the *Drosophila* Secretome database.

Genes predicted as cell surface proteins were retrieved from two publications and used to supplement *Drosophila* Secretome (Kurusu *et al*. 2008; Pei *et al*. 2018). Genes identified in exosome vesicles were retrieved from Vesiclepedia database (Kalra *et al*. 2012; Pathan *et al*. 2019; Chitti *et al*. 2024) (https://www.microvesicles.org/), then the subset of genes on this list from *Drosophila melanogaster* were selected and integrated into the *Drosophila* Secretome database.

### Bioinformatics analysis of *Drosophila* secretome

DIOPT vs10 (https://www.flyrnai.org/diopt) was used to analyze the conservation of *Drosophila* secretome (Figure 1B). A rank filter was used to exclude low confidence mapping (Hu *et al*. 2011; Hu *et al*. 2026). The human Secretome was assembled using the secretome annotation downloaded from the Human Protein Atlas (June 3^rd^ 2026) (https://www.proteinatlas.org/download/proteinatlas.tsv.zip) (Uhlen *et al*. 2019) and several secreted gene databases including SPD, SPRomeDB, and SEPDB (Chen *et al*. 2005; Chen *et al*. 2019; Wang *et al*. 2024). Mouse and rat secretomes were obtained from SEPDB (Wang *et al*. 2024) and a zebrafish secretome was obtained from publication (Klee 2008).

PANGEA (Hu *et al*. 2023) was used for Gene Set Enrichment Analysis with a significance threshold of *p*-value < 0.05 with the generic gene ontology SLIM terms for “biological process” was selected. A bar graph displaying the enrichment *p*-value and log2 fold changes was downloaded from PANGEA (Figure 1C). Secretome genes were further categorized based on the enrichment results. For example, genes involved in an immune system process, nervous system process, behavior, or development were annotated based on gene ontology annotation obtained from FlyBase (FB2026_3), and genes categorized as metabolic enzymes were annotated based on the intersection of the metabolic process by gene ontology and enzyme annotation by FlyBase Gene Group annotations.

Tissue specificity analysis was performed using the integrated Fly Cell Atlas (FCA) dataset (Li *et al*. 2022). Specifically, the “10x VSN All (Stringent)” dataset was downloaded as a loom file from https://www.flycellatlas.org/#data and the 250 annotated cell types within FCA were manually consolidated into 14 biological systems (Sup. Table 1). To obtain robust, system-level expression estimates, a pseudo-bulking strategy was applied: raw UMI counts from all cells belonging to the same system were summed to produce a single aggregated count vector per system, which was then normalized to counts per million (CPM) to account for differences in cell number and sequencing depth across systems. Each gene was subsequently assigned to one of six tissue specificity categories (“Tissue Enriched”, “Group Enriched”, “Tissue Enhanced”, “Mixed”, “Expressed in All”, and “Not Detected”), following the classification framework established by the Human Protein Atlas (Karlsson *et al*. 2021) using a 5-fold expression difference as the threshold. A gene is classified as “Tissue Enriched” if its CPM in one system is at least 5-fold higher than in any other system; “Tissue Enhanced” if its CPM in one system is at least 5-fold higher than the average CPM of all other systems without meeting the stricter tissue-enriched criterion; “Group Enriched” if, in a group of two to five systems, all CPM values within the defined group are at least 5-fold higher than all other systems. For genes that do not meet the criteria for the enriched or enhanced classifications, those with moderate expression across all systems (all CPM > 10) are classified as “Expressed in All”, those without detectable expression across all systems (all CPM < 10) are “Not Detected”, and all remaining genes are classified as “Mixed” (Figure 2A). The secreted genes from the high rank tier were used for hierarchical clustering analysis using the gene expression matrix of CPM values. Log10 (CPM+1) values were used to represent the expression levels. A heatmap visualization was created using the ComplexHeatmap (v2.26.1) package in R, clustering both rows and columns by Euclidean distance and adding annotation bars to show the characterization of the secreted genes (Figure 2C).

**Figure 2.**
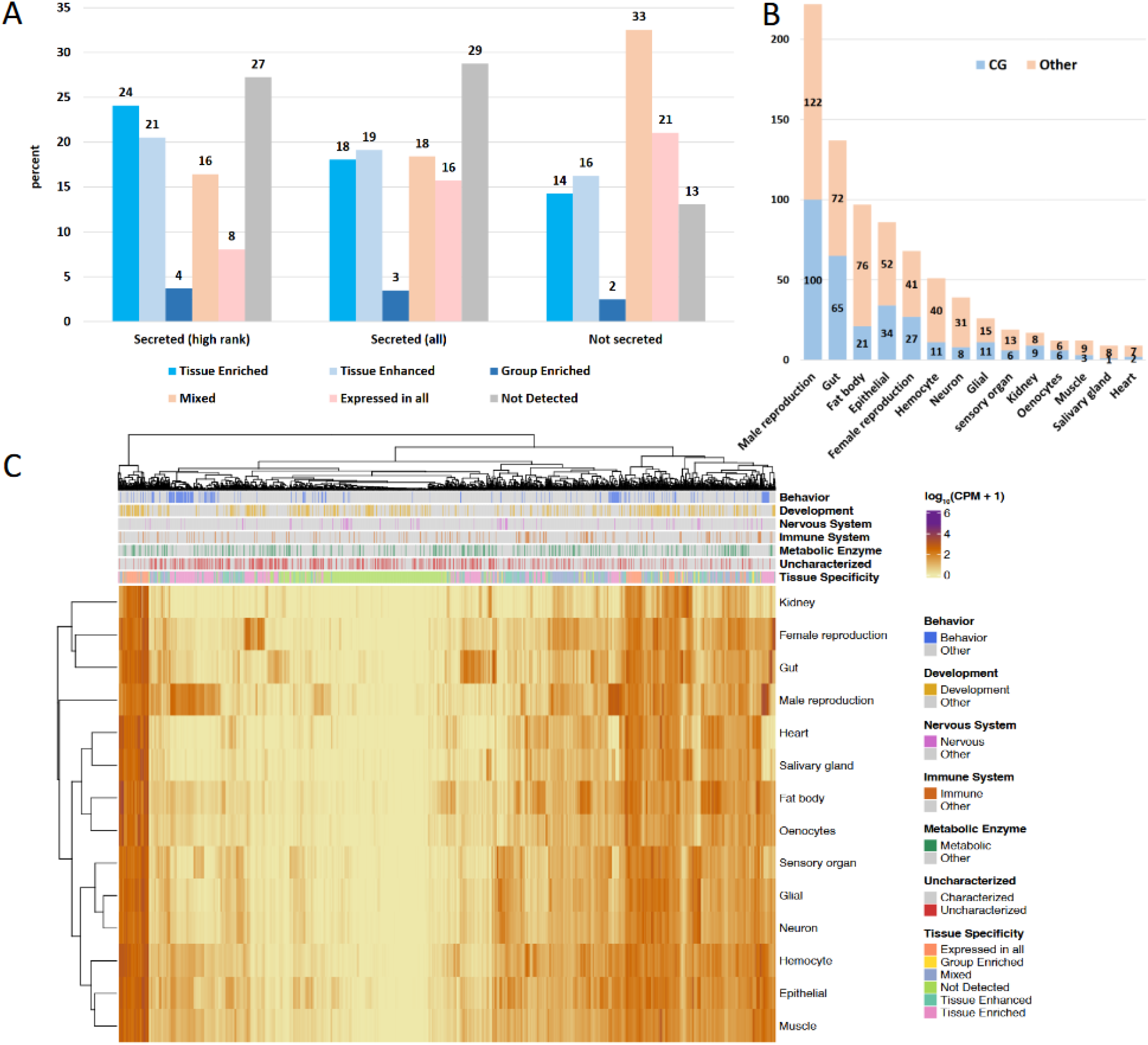
Tissue specificity analysis. A.) Statistics of tissue expression group B.) Tissue enriched gene distribution. A.) Clustering analysis of ligands (high-confident rank) expression based on the snRNA-seq datasets from Fly Cell Atlas. The 250 cell types were manually grouped into 14 broad systems and pseudo-bulk expression profiles were generated by summing raw UMI counts across all cells of the same annotated system within each biological sample, followed by CPM normalization to produce a single representative expression estimate per cell type per gene.

The Aging Fly Cell Atlas (AFCA) dataset was obtained from (Lu *et al*. 2023) and accessed from NCBI GEO under the accession GSE218661. Only the body data was utilized, excluding the head. The raw counts, normalized counts, and metadata were downloaded and used to create a Seurat object that was used for the remainder of the downstream analysis. Specific cell types were manually mapped to systems using the FCA and AFCA annotations (Sup. Table 1). Any cells left unannotated or with conflicting FCA and AFCA annotations were removed from further analysis. The aging atlas data was pseudo-bulked by system at each age (5-day, 30-day, 50-day, and 70-day) in R (v4.5.0). The Seurat object was subsetted to the age of interest, and the raw counts matrix was extracted. The sum of gene counts in all cells belonging to each system was taken to pseudobulk the data, and each system’s library size was normalized by CPM. Differentially expressed gene (DEG) calling was performed using the Seurat object prepared as previously described and following a standard Seurat workflow (v5.5.0). For each condition tested, DEGs were calculated per cell type using the older age as the experimental condition using Seurat’s FindMarkers function. They were then filtered down to results with an adjusted *p*-value < 0.05 and absolute log2 fold change > 1. The secreted genes that were also found to be DEGs when comparing 70-day old flies vs 5-day old flies were selected for hierarchical clustering analysis. Similar to the FCA clustering, a heatmap of tissues vs secreted DEGs was visualized using the log2 fold change values to illustrate changes in expression. A heatmap visualization was created using the ComplexHeatmap (v2.26.1) package in R, clustering both rows and columns by Euclidean distance and adding annotation bars to show the characterization of the secreted genes (Figure 4C).

### Implementation of the online portal

The *Drosophila* Secretome web application (www.flyrnai.org/apps/fly_secretome/) was built with the Flask web framework and is served by the Gunicorn WSGI server. The underlying data for the application is stored in a MySQL database, which is accessed via SQLAlchemy. Python is used for the backend of the application, while the frontend uses HTML templates rendered with Jinja. JavaScript is used throughout for AJAX requests and DOM manipulation. Bootstrap is used for layout and styling, and Font Awesome for project iconography. Both the web application and its database are hosted on the O2 high⍰performance computing (HPC) cluster at Harvard Medical School, operated by the Research Computing group.

## RESULTS

### Assembly of a *Drosophila* secretome database

To assemble the candidate gene list of putative *Drosophila* secretome, we collected genes annotated from multiple resources first: FlyBase (Ozturk-Colak *et al*. 2024), UniProt (UniProt 2023), Gene Ontology (Gene Ontology 2026), and a recent large-scale secretome dataset(Bosch *et al*. 2026). FlyBase annotates gene products that interact with a receptor to modulate its activity, curated from the primary literature. UniProt annotates secreted proteins as part of its subcellular localization schema, and the Gene Ontology Consortium annotates gene products localized to the extracellular region under the Cellular Component ontology. The fourth source is a recent study (Bosch *et al*. 2026) that presents a comprehensive map of the *Drosophila* secretome using TurboID proximity labeling targeted to the ER lumen, combined with large-scale hemolymph isolation and quantitative mass spectrometry, identifying 535 circulating proteins secreted from 10 major larval tissues. Together, these four sources yielded 1802 unique genes. Of these, 1,130 (63%) were contributed by a single source, while 672 (37%) appeared in two or more sources (Figure 1A).

Since annotation and data coverage for secreted proteins remains incomplete, we complemented the curated gene list with proteome-scale computational predictions. Signal peptide prediction tools including SignalP (Teufel *et al*. 2022), TargetP (Emanuelsson *et al*. 2000; Emanuelsson *et al*. 2007), Phobius (Kall *et al*. 2007) and TOPCONS (Tsirigos *et al*. 2015) and a subcellular localization predictor (DeepLoc2) (Nielsen 2025) were applied across the entire *Drosophila* proteome. The predictions showed substantial agreement with the annotated set: 1,569 of the 1,802 curated genes (87%) had at least one protein isoform with a signal peptide predicted. At the same time, 2,509 additional genes not covered by any of the four annotation sources were computationally predicted to encode signal peptides and may represent bona fide secreted proteins. To incorporate these candidates while reflecting their lower evidential support, we expanded the secretome database to include the predicted set but assigned it a lower confidence tier. The 1,802 genes derived from curated annotation resources and experimental data were retained at the high-confidence tier.

Several key signaling pathways rely on the proteolytic processing of membrane-anchored proteins as a mechanism for releasing extracellular signals. A well-characterized example is the EGFR signaling pathway in *Drosophila melanogaster*, in which the primary ligand (Spitz) is synthesized as an inactive transmembrane precursor. Then Star binds Spitz and escorts it out of the ER into the Golgi where the intramembrane serine protease Rhomboid-1 (Rho-1) cleaves Spitz within its transmembrane domain. Cleavage releases the N-terminal EGF domain of Spitz as a soluble, diffusible ligand, which is secreted from the cell and can travel to activate the EGFR signaling pathway on neighboring or distant cells (Lee *et al*. 2001; Urban *et al*. 2001). To further expand the database to include this class of conditionally secreted proteins, we incorporated the genes predicted as cell surface proteins from multiple publications of the large-scale prediction effort (Kurusu *et al*. 2008; Pei *et al*. 2018). Furthermore, we applied transmembrane helix prediction algorithms, such as deepTMHMM (Hallgren *et al*. 2022) alongside ProP (Duckert *et al*. 2004), a tool for proteolytic cleavage site prediction. The results of these analyses are integrated into the *Drosophila* Secretome database as an additional annotation layer, providing users with putative candidates to support hypothesis generation and experimental design in contexts where regulated ectodomain shedding or intramembrane proteolysis may be relevant.

In addition, we also incorporated genes whose products are transported out of the cell through exosome-mediated release, in which cells package selected proteins, lipids, and nucleic acids into small extracellular vesicles that serve as vehicles for intercellular communication. To capture this class of secreted proteins, we mined the Vesiclepedia database (Chitti *et al*. 2024), which catalogs proteins identified in extracellular vesicles across multiple organisms including *Drosophila*. This added 451 genes not previously included in the database, further broadening the coverage of the *Drosophila* secretome beyond classically secreted proteins.

In total there are 5,280 protein-coding genes in the *Drosophila* secretome we assembled (Sup. Table 2A, Sup Figure 1A). Of these, 1,802 are based on annotation and were assigned to the high-confidence tier (Figure 1A), and 3,478 genes are based on computational predictions or inclusion in Vesiclepedia, and were assigned to the low-confidence tier. Besides rank assignment for confidence tier, a confidence score is also assigned for every gene. For example, a gene annotated in both UniProt and GO receives a higher score than the one annotated in only one of them, and a gene supported by direct experimental evidence receives a higher score than one supported by other types of evidence, such as homology-based evidence (Sup. Table 2B).

### Bioinformatics analysis of *Drosophila* secretome

#### 1.) Conservation

Using the ortholog mapping tool DIOPT (v10; confidence filter: high or moderate) (Hu *et al*. 2011; Hu *et al*. 2026), approximately 70% of *Drosophila* genes have a human ortholog. Interestingly, secretome genes are less conserved than the rest of the genome with only 56% of the *Drosophila* secretome (2,955 out of 5,280 genes) having a human ortholog, compared to 79% of non-secretome *Drosophila* genes. This suggests that a large fraction of secreted proteins is likely to serve species-specific functions. Regarding the portion conserved with human, we compared it with human secretome annotation. We first assembled a reference list of 6881 human secretome genes by integrating candidates from the secretome annotation of Human Protein Atlas (HPA) (Uhlen *et al*. 2019) and three dedicated secreted protein databases: SPD, SPRomeDB, and SEPDB (Chen *et al*. 2005; Chen *et al*. 2019; Wang *et al*. 2024). Of the 2,955 conserved *Drosophila* secretome genes, the human orthologs of 2,402 (81%) are independently annotated as human secretome genes (Figure 1B). This strong cross-species concordance supports the utility of the fly as a model for studying conserved extracellular signaling.

We also analyzed the length of the longest isoform for each *Drosophila* gene and found that 28% of secreted genes encode proteins shorter than 200 amino acids, compared to only 19% in the rest of the genome. Short proteins yield short pairwise alignments with lower bit scores and E-values even for true orthologs, making it harder for alignment-score-dependent pipelines to distinguish orthologs from spurious hits or paralogs; they also carry less phylogenetic signal for tree-based reconciliation and are more prone to partial or domain-only alignments that confound reciprocal-best-hit approaches (Trachana *et al*. 2011). To evaluate whether the overall conservation gap seen between secreted and non-secreted genes is due to the technical limitation of short-sequence alignment, we analyzed conservation while accounting for protein length by separating the genes into two groups: those encoding proteins longer than 200 amino acids and those encoding proteins 200 amino acids or shorter, and then calculated the proportion conserved in each group (Sup. Figure 1B). The gap in conservation between secreted and non-secreted genes was modest but still present for larger proteins (69% vs. 83%) and much larger for smaller proteins (31% vs. 60%). This analysis indicates that size does have a relationship with identification of orthologs but even after taking protein size into account, there is still a conservation gap between secreted and non-secreted genes.

### Biological process

Gene set enrichment analysis (GSEA) was done on the subset of high rank *Drosophila* secretome using PANGEA (Hu *et al*. 2023) based on the generic gene ontology slim terms of biological process. The analysis revealed significant enrichment for genes involved in immune defense, metabolic regulation and nervous system process (Figure 1C), which is consistent with the established roles of secreted proteins as key mediators of cell-cell signaling and inter-organ communication. For example, the immune system relies heavily on circulating cytokines and antimicrobial peptides (Ferrandon *et al*. 2007); the nervous system employs secreted neuropeptides and growth factors for synaptic modulation and axon guidance (Ballard *et al*. 2014; Onesto *et al*. 2021); and metabolic homeostasis is coordinated in large part through hormonally active circulating factors such as insulin-like peptides and adipokines (Ahmad *et al*. 2020).

#### 2.) Tissue expression

The Fly Cell Atlas (Li *et al*. 2022) is a comprehensive single-nucleus RNA-seq resource for *Drosophila*. The underlying dataset profiles 580,000 nuclei from 15 individually dissected tissues of both male and female adult flies as well as the entire head and body, annotated to over 250 distinct cell types. The annotations were contributed by a consortium effort from over 100 experts of 40 international *Drosophila* research laboratories, and the dataset serves as a community reference for gene expression pattern in wild type adult flies. To summarize expression patterns at a higher level of biological organization, we grouped 250 annotated cell types into 14 broad systems (Sup. Table 1). For example, all glial cell types were consolidated into a single “glia” category, and all neuronal subtypes into a single “neuron” category. Pseudo-bulk expression profiles were then generated for each system by aggregating raw UMI counts across all constituent cells, and expression levels were normalized to counts per million (CPM), yielding a single comparable expression estimate per gene per system.

Tissue specificity analysis was done using an established method (Karlsson *et al*. 2021) to group genes into 6 categories: “Expressed in all,” “Tissue Enriched,” “Group Enriched,” “Tissue Enhanced,” “Mixed,” and “Not Detected”. The secretome has a higher portion of tissue-specific genes and smaller portion of universally expressed genes than the rest of the proteome, indicating that secreted genes are less likely to be housekeeping genes and more likely to play tissue-specific roles (Figure 2A). For example, among the tissue enriched genes, many genes are specifically expressed in reproductive organs (Figure 2B). To visualize the expression landscape of the secretome, we performed hierarchical clustering on the CPM values of 1,802 genes from the high-confidence tier and displayed the result using a heatmap along with annotations of tissue specificity and various biological processes (Figure 2C). The majority of ligands show moderate to low expression (light yellow to orange) across most systems, with a relatively small subset showing high expression (dark brown to purple) in a tissue-restricted manner. This sparse high-expression pattern is consistent with the biology of secreted ligands, which tend to be produced in specific source tissues rather than ubiquitously. Moreover, in the row dendrogram, reproductive and excretory tissues (kidney, female reproduction, male reproduction) cluster together, suggesting shared ligand expression programs, while neural and immune tissues (glia, neuron, hemocyte) form a loose grouping, as do metabolically active tissues (fat body, oenocytes, muscle). The fat body and gut rows show notably broader and higher expression across many ligand columns, consistent with their roles as major secretory organs that produce a large proportion of circulating factors as also confirmed by the secretome proteomics data (Bosch *et al*. 2026). The immune-annotated ligands appear scattered, but there is some clustering consistent with the finding that hemocytes and the fat body are the primary immune secretory sources. Additionally, nervous system ligands cluster in specific column groups, reflecting the specialized signaling repertoire of neurons and glia. To further illustrate the distribution of tissue specifically expressing ligands over various biological process, we calculated the ratio of genes annotated in each biological process among all genes specifically expressing in each tissue, then displayed the results as a heatmap (Sup. Figure 3). Overall these results demonstrate that part of the *Drosophila* secretome is functionally organized in a highly tissue-specific manner, with each tissue contributing a distinct subset of secreted ligands aligned with its primary biological function.

There are 5,834 uncharacterized protein-coding genes in *Drosophila* (FB2026_3) which makes up 42% of the *Drosophila* proteome. We noticed that there are slightly more uncharacterized genes in the *Drosophila* secretome (44%) compared to the remaining genes (40%). In addition, 48% of genes included in the secretome based only on computational predictions are CG genes. The comparison of the FCA expression between the uncharacterized and characterized genes of *Drosophila* secretome (Sup. Figure 2) demonstrated that among uncharacterized genes there are more undetected genes (40% vs 18%) and fewer universally expressed genes (4% vs 24%). There are also more tissue-specific genes expressed among CG genes (22% vs 14%), which suggests an opportunity for functional discovery in a tissue-specific manner (Figure 2B). *Drosophila* has one of the richest resources of genetic reagents among model organisms, largely due to decades of community-driven stock center investment (Mohr *et al*. 2014; Bilder and Irvine 2017). VDRC (Vienna *Drosophila* Resource Center), TRiP (Transgenic RNAi Project) and NIG (National Institute of Genetics) (Dietzl *et al*. 2007; Perkins *et al*. 2015; Zirin *et al*. 2020) together provide RNAi transgenic lines covering 92% of secretome genes, including 85% of the 2,208 uncharacterized secretome genes. Additionally, 33% of the secretome and 15% of the uncharacterized secretome are covered by UAS-fly-cDNA or UAS-human-cDNA stock collection (Bischof *et al*. 2013; Kanca *et al*. 2022; Avila *et al*. 2024) that could be used for gain-of-function experiments. The combination of transgenic stocks and tissue-specific driver lines makes it possible to move the large uncharacterized fraction of the secretome identified here efficiently from computational prediction to functional testing.

### *Drosophila* Secretome online resource

The *Drosophila* Secretome online resource provides a dedicated portal for researchers to explore and mine the database (Figure 3). Users can search a single gene to determine whether it is part of the secretome. If so, the portal retrieves and displays the evidence source, confidence tier, signal peptide predictions and subcellular localization predictions. In addition, each entry is cross-referenced against available proteomics datasets, allowing users to determine whether the protein has been detected in the larval circulatory system and, if so, its tissues of origin.

**Figure 3.**
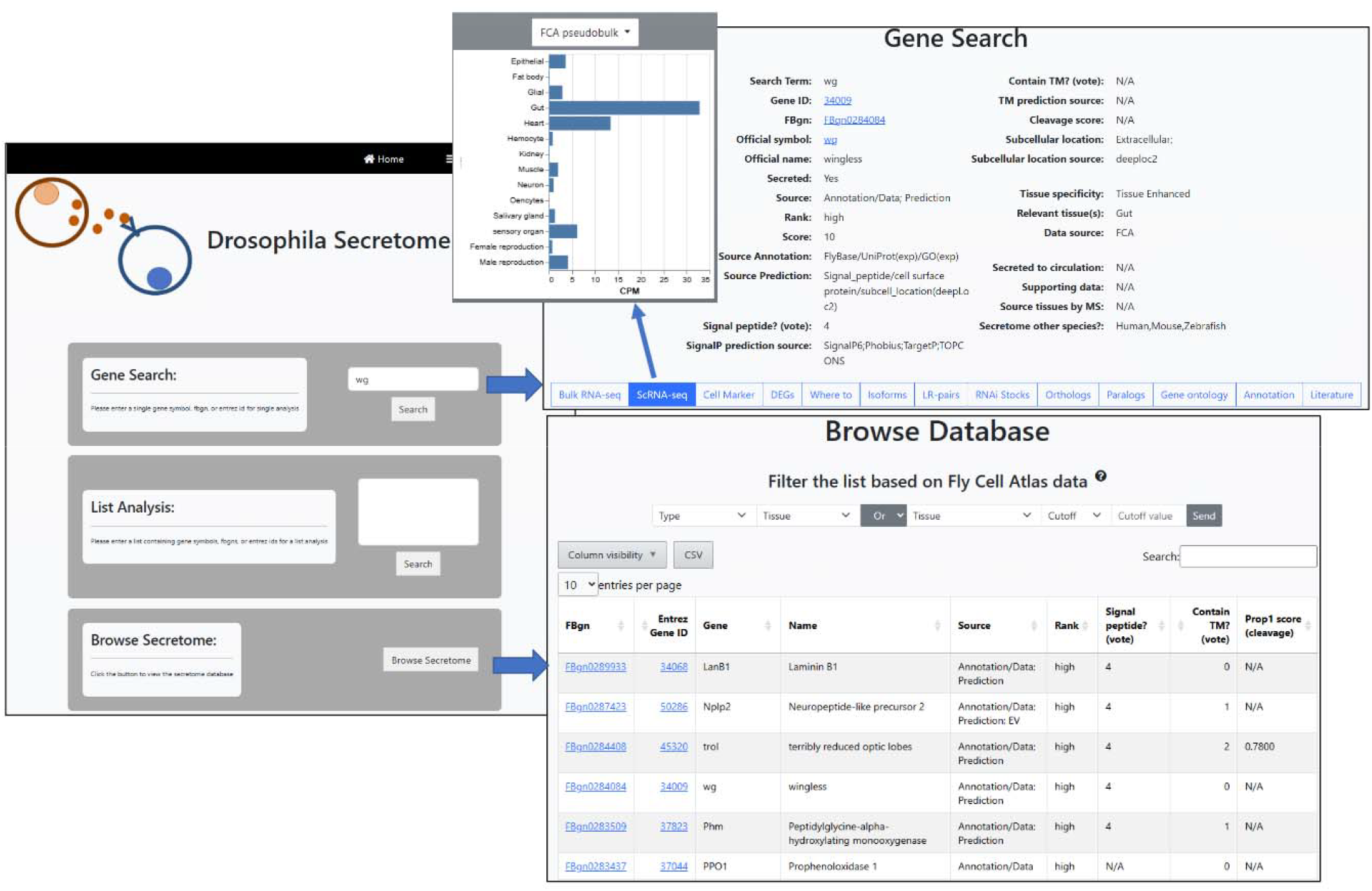
*Drosophila* Secretome online portal was developed for user to do customized search. User can perform one gene search, and mine the data, annotation, literature as well as transgenic fly lines associated with the input gene. User can also browse the Secretome database and apply expression filter based on the snRNA-seq datasets from Fly Cell Atlas.

A menu bar with 13 tabs provides structured access to additional information relevant to experimental design (Figure 3 and Sup. Figure 4). For transcriptomic analysis, a dedicated tab enables mining of bulk RNA-seq datasets, including tissue, cell line, developmental stage, and treatment-focused datasets from the modENCODE (Boley *et al*. 2014) and FlyAtlas2 consortia (Krause *et al*. 2022), gut subregion and cell-type transcriptomic profiles (Marianes and Spradling 2013; Dutta *et al*. 2015), primary cell line transcriptomes (Coleman-Gosser *et al*. 2023), and selected time-course datasets from our lab (Zirin *et al*. 2020; Saavedra *et al*. 2023). Single-cell RNA-seq data are accessible through three separate tabs covering: (1) pseudo-bulk expression values from two full body datasets including FCA dataset (Li *et al*. 2022; Liu *et al*. 2025); (2) cell-type-specific marker genes; and (3) differential expression across genotypes or treatment conditions, drawn from all published scRNA-seq datasets from our lab (Petsakou *et al*. 2023; Saavedra *et al*. 2023; Xu *et al*. 2024; Li *et al*. 2025; Liu *et al*. 2025; Ewen-Campen *et al*. 2026; Lane *et al*. 2026). One tab is dedicated to answer where the protein is secreted to eg. circulating system or exosome vesicle based on proteomics data (Kalra *et al*. 2012; Pathan *et al*. 2019; Chitti *et al*. 2024; Bosch *et al*. 2026). Additional tabs provide ortholog and paralog predictions from DIOPT (Hu *et al*. 2011; Hu *et al*. 2026), gene set annotations from PANGEA (Hu *et al*. 2023), candidate receptors annotated in FlyPhoneDB (Liu *et al*. 2022; Qadiri *et al*. 2025) or predicted by FlyPredictome (Kim *et al*. 2026), and RNAi stock information to facilitate the identification of transgenic fly lines for functional follow-up.

Beyond single-gene queries, users can perform batch searches by submitting a gene list to identify the secreted subset along with the associated annotations. Users can also browse the complete secretome and apply filters to select candidate secreted genes expressed in one or more tissues of interest, based on FCA transcriptomic data.

### Global expression changes in the secretome during aging in *Drosophila*

The Aging Fly Cell Atlas (AFCA) is a large-scale single-cell transcriptomics study designed to characterize how aging affects gene expression, cell composition, and cellular states across the entire adult *Drosophila melanogaster* body (Lu *et al*. 2023). By profiling hundreds of cell types at multiple ages, the study provides a comprehensive resource for identifying cell type–specific signatures of aging, including changes in metabolism, stress responses, immune activity, and tissue homeostasis. We reformed the data from young adult flies (5 day) and old adult flies (70 day) to higher order tissue groups then performed differential gene expression analysis by systems. This analysis revealed that secreted genes are more likely to be differentially regulated than non-secreted genes (Figure 4A). Specifically, 39% of the secreted genes from the high confident tier are down-regulated in one or multiple tissues during aging compared to 22% of non-secreted genes. We further examined the distribution of secreted genes over the source tissues, and the female reproduction system and gut have the largest number of down-regulated secreted genes (Figure 4B). Across almost all tissues, the number of secreted genes that are down-regulated with age far exceeds those that are up-regulated, often by 5–15 fold. Female reproduction, gut, and fat body, tissues which are central to reproduction, nutrient sensing, and systemic metabolic signaling, respectively, show the steepest down-regulation of secreted genes, consistent with known age-related declines in fertility and metabolic homeostasis in *Drosophila*. In contrast, neuron and sensory organ show smallest absolute numbers of up-regulated genes (~5 each) and a moderate number of down-regulated genes, suggesting these post-mitotic tissues have a more limited but still declining secretory transcriptional response with age. In addition, we also evaluated the gene groups among the DEGs in each tissue (Figure 4C, D) and confirmed that, as in the gut, many metabolic enzymes are down-regulated. As flies get older, their innate immune pathways become progressively and inappropriately active even in the absence of infection; indeed, this is one of the most reproducible molecular signatures of aging flies. In hemocytes, even though there are more down-regulated genes than up-regulated genes overall, more genes involved in immune system are up-regulated than down-regulated, which is consistent with the findings that chronic immune activation occurs during aging. Overall, our analysis confirmed the classic aging signature of broad transcriptional decline in metabolic, developmental, and structural/behavioral gene programs across nearly all tissues and dysregulation of immune-related genes in both directions. Together, these results position secreted factors not merely as passive readouts of tissue aging but as candidate mediators of aging.

**Figure 4.**
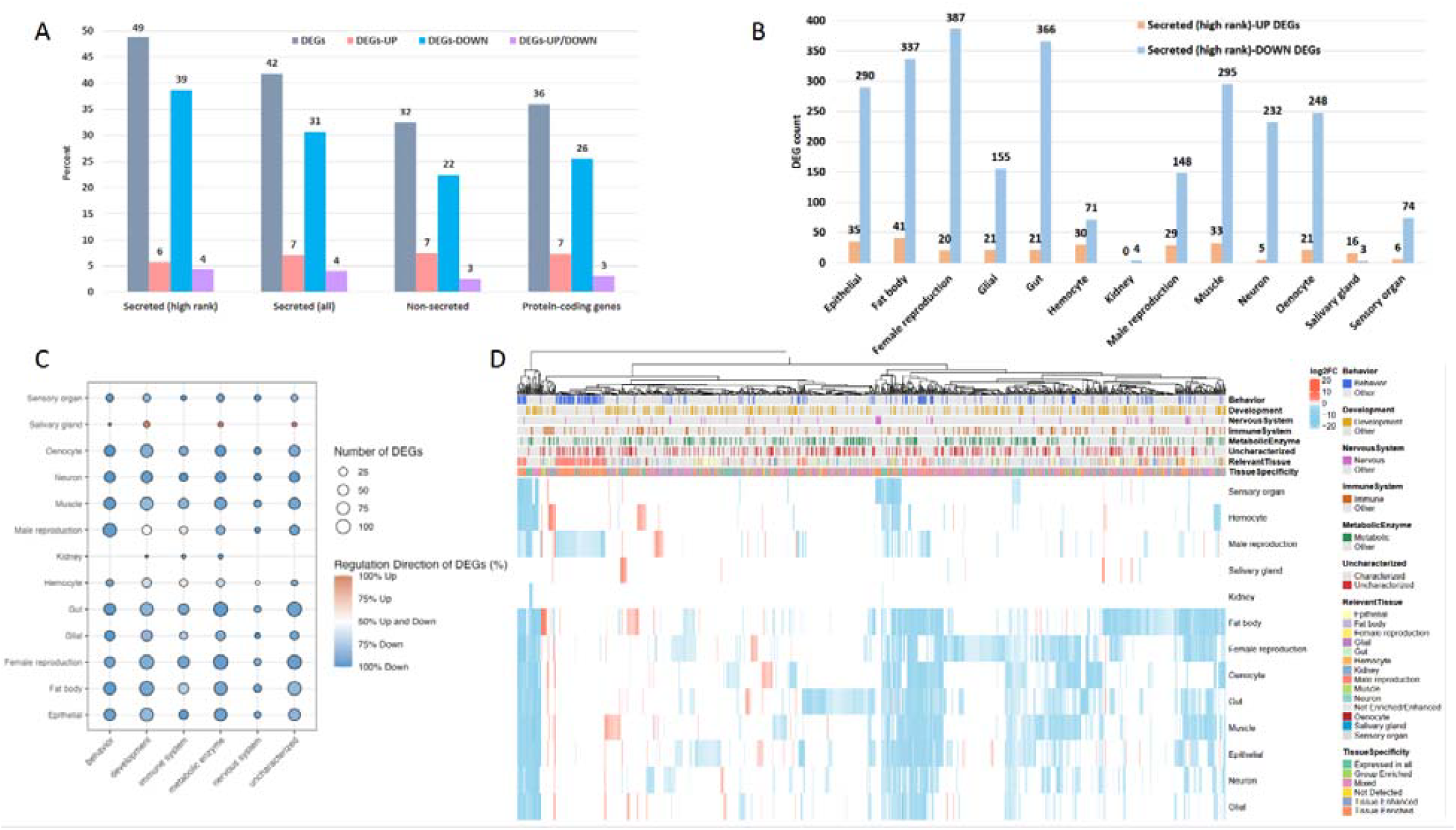
Secretome changes during aging in Drosophila. A.) Differentially expressed genes were identified comparing aged fly (70 day) and young adult fly (5 day) using AFCA data. With the cutoff of *p*-value adj < 0.05 and abs(log2fc)>1, 36% of the protein coding genes are either up-regulated or down-regulated in one or more tissues while 49% of secreted genes in high confidence tier are DEGs. B.) Counts of high-confident secreted DEGs from various tissue types. C.) The distribution of various gene groups among secreted DEGs in each tissue. D.) Hierarchical clustering of secreted DEGs using the criteria of adjusted P value <0.05 and log2 fold change >1 or <-1.

## DISCUSSION

Here, we present a comprehensive database of the *Drosophila* secretome that integrates curated annotation and proteome-wide computational prediction for conventionally secreted proteins, as well as information for unconventional secretion pathway proteins, and characterize the evolutionary, functional, and tissue-specific properties of secreted proteins in the fly. By combining four independent annotation sources with signal peptide, subcellular localization predictions, and extracellular vesicle proteomics, we expand the annotated *Drosophila* secretome from 1,802 to 5,280 candidate genes while the rank annotation distinguishes the genes with strong experimental or curatorial support from those nominated purely by sequence-based prediction, allowing the resource to serve both as a high-confidence reference set and as a broader discovery tool for genes whose secretion has not yet been experimentally validated.

Our finding that only 56% of *Drosophila* secretome genes have a human ortholog, compared to 79% of the remainder of the genome, indicates that secreted proteins are markedly less conserved than the proteome as a whole. This is consistent with the general expectation that extracellular signaling molecules, particularly those mediating immune defense, chemo sensing, and reproduction, are subject to more rapid evolutionary turnover than intracellular housekeeping machinery, likely reflecting species-specific ecological pressures such as pathogen exposure, mating systems, and environmental adaptation. At the same time, among the conserved subset, 81% concordance with independently annotated human secretome genes reinforces the utility of *Drosophila* as a model for studying the more deeply conserved part of extracellular signaling biology.

The enrichment of tissue-restricted expression among secretome genes, relative to the higher proportion of ubiquitously expressed housekeeping genes in the remaining proteome, supports a model in which secreted ligands are produced by specialized source tissues to coordinate inter-organ communication, rather than being broadly and constitutively expressed. The clustering of reproductive and excretory tissues, and separately of metabolically active tissues such as fat body and gut, aligns with known physiological roles of these organs as major endocrine and metabolic signaling hubs, and is corroborated by independent proteomic detection of circulating factors originating from these tissues.

The modest enrichment of uncharacterized (CG) genes within the secretome, and particularly their over-representation among computationally predicted, low-confidence candidates, points to a substantial blind spot in current functional annotation. The observation that CG genes show more tissue-restricted expression than characterized secretome genes, despite being annotated at lower confidence, suggests that at least some of these predicted secreted proteins may have genuine, tissue-specific signaling roles that have simply not yet been investigated. This positions the uncharacterized fraction of the secretome as a promising target for hypothesis-driven functional studies, particularly using the tissue-expression filters built into the accompanying web resource.

The disproportionate down-regulation of secreted genes relative to the rest of the transcriptome during aging, particularly in the gut, fat body, and female reproductive system, is consistent with established models in which age-related decline in inter-organ communication contributes to loss of metabolic and reproductive homeostasis. In contrast, the pattern in hemocytes where secreted immune genes were preferentially up-regulated despite an overall decline in secreted gene expression, reinforces the well-documented phenomenon of age-associated chronic innate immune activation in the absence of infection, and highlights that the secretome does not decline uniformly but is instead selectively dysregulated in a biological process and tissue-dependent manner.

Several caveats merit consideration: First, the low-confidence tier of the database that relies on computational prediction of signal peptides, makes the significant portion of the resource and has not been experimentally validated, therefore, false positives and false negatives are expected at rates inherent to these prediction tools. In addition, proteins in the secretome fall into three categories: blood proteins, locally secreted proteins, and intracellular proteins (Uhlen *et al*. 2019). The intracellular group enters the ER but is then routed to other compartments (such as lysosomes) or retained in the ER or Golgi, so these proteins are never secreted from the cell. They were identified using DeepLoc2 subcellular localization predictions, assigned a low score, and flagged so that users can easily filter them out, since they are of less interest for studies of organ-to-organ communication. Given the inherent limitations of subcellular localization prediction, some proteins in this group may be misclassified. To further improve the annotation, a feedback page is implemented at the online portal and further annotation by the community will help refine the resource over time. Second, the tissue expression analyses are derived from pseudo-bulk aggregation of single-nucleus data, which may obscure cell-type-specific expression patterns within the broader tissue categories used here. Third, cross-species conservation analysis was restricted to sequence-based ortholog, therefore, some diverged signaling molecules may have been missed. Future integration of structural search algorithms, such as Foldseek (van Kempen *et al*. 2024), could overcome this limitation by uncovering remote homologs and structurally conserved factors.

As summary, this work provides both a comprehensive resource and a set of biological insights into the organization, evolution, and physiological relevance of the *Drosophila* secretome. The online resource with the integration of transcriptomic, proteomic, and functional genetic data offers a framework for prioritizing candidates for future study, particularly among the large uncharacterized fraction of the secretome. The convergence of transcriptional aging signatures and functional longevity data in the gut secretome suggests that systematic, tissue-directed screening of secreted factors may reveal new mediators of inter-organ communication relevant to aging and disease, with potential relevance to conserved human orthologs identified through this same framework.

## ACKNOWLEDGEMENTS

We thank the past and current members from Perrimon lab for their valuable suggestions and feedbacks. Particularly we would like to thank Dr. Stephanie Mohr for help with paper writing. We extend our gratitude to the Harvard Medical School Research Computing and IT-Client Services teams for their consultation, web hosting, and support.

## FUNDING

This work was supported in part by a grant from the U.S. National Institutes of Health (NIH) National Institute of General Medical Sciences (P41 GM132087) as well as by grants from the NIH Office for Research Infrastructure Projects (R24 OD026435, R24 OD030002, R24 OD019847, R24 OD031952) to support resource development. N.P. is an investigator of Howard Hughes Medical Institute.

## DATA AVAILABILITY STATEMENT

The resources are available to pubic. The URL for *Drosophila* secretome database is https://www.flyrnai.org/apps/fly_secretome.

## CONFLICTS OF INTERESTS

The authors declare no conflicts of interest.

## Notes

https://www.flyrnai.org/apps/fly_secretome

### Competing Interest Statement

The authors have declared no competing interest.

### Summary of Updates

The manuscript was extensively revised by the authors and supplemental material has been added.

## REFERENCES

Ahmad, M., L. He and N. Perrimon, 2020 Regulation of insulin and adipokinetic hormone/glucagon production in flies. Wiley Interdiscip Rev Dev Biol 9: e360.

Avila, A., L. Paculis, R. G. Tascon, B. Ramos and D. Jia, 2024 A large-scale in vivo screen to investigate the roles of human genes in Drosophila melanogaster. G3 (Bethesda) 14.

Ballard, S. L., D. L. Miller and B. Ganetzky, 2014 Retrograde neurotrophin signaling through Tollo regulates synaptic growth in Drosophila. J Cell Biol 204: 1157–1172.

Bilder, D., and K. D. Irvine, 2017 Taking Stock of the Drosophila Research Ecosystem. Genetics 206: 1227–1236.

Bischof, J., M. Bjorklund, E. Furger, C. Schertel, J. Taipale et al., 2013 A versatile platform for creating a comprehensive UAS-ORFeome library in Drosophila. Development 140: 2434–2442.

Boley, N., K. H. Wan, P. J. Bickel and S. E. Celniker, 2014 Navigating and mining modENCODE data. Methods 68: 38–47.

Bosch, J. A., P. M. J. Beltran, C. Cavers, J. T. LaGraff, R. Melanson et al., 2026 Multi-omic mapping of Drosophila protein secretomes reveals tissue-specific origins and inter-organ trafficking. Nat Commun 17.

Chen, G., J. Chen, H. Liu, S. Chen, Y. Zhang et al., 2019 Comprehensive Identification and Characterization of Human Secretome Based on Integrative Proteomic and Transcriptomic Data. Front Cell Dev Biol 7: 299.

Chen, Y., Y. Zhang, Y. Yin, G. Gao, S. Li et al., 2005 SPD--a web-based secreted protein database. Nucleic Acids Res 33: D169–173.

Chitti, S. V., S. Gummadi, T. Kang, S. Shahi, A. L. Marzan et al., 2024 Vesiclepedia 2024:an extracellular vesicles and extracellular particles repository. Nucleic Acids Res 52: D1694–D1698.

Coleman-Gosser, N., Y. Hu, S. Raghuvanshi, S. Stitzinger, W. Chen et al., 2023 Continuous muscle, glial, epithelial, neuronal, and hemocyte cell lines for Drosophila research. Elife 12.

Dietzl, G., D. Chen, F. Schnorrer, K. C. Su, Y. Barinova et al., 2007 A genome-wide transgenic RNAi library for conditional gene inactivation in Drosophila. Nature 448: 151–156.

Duckert, P., S. Brunak and N. Blom, 2004 Prediction of proprotein convertase cleavage sites. Protein Eng Des Sel 17: 107–112.

Dutta, D., A. J. Dobson, P. L. Houtz, C. Glasser, J. Revah et al., 2015 Regional Cell-Specific Transcriptome Mapping Reveals Regulatory Complexity in the Adult Drosophila Midgut. Cell Rep 12: 346–358.

Emanuelsson, O., S. Brunak, G. von Heijne and H. Nielsen, 2007 Locating proteins in the cell using TargetP, SignalP and related tools. Nat Protoc 2: 953–971.

Emanuelsson, O., H. Nielsen, S. Brunak and G. von Heijne, 2000 Predicting subcellular localization of proteins based on their N-terminal amino acid sequence. J Mol Biol 300: 1005–1016.

Ewen-Campen, B., W. Chen, S. G. Tattikota, Y. Liu, Y. Hu et al., 2026 The Drosophila proventriculus lacks stem cells but compensates for age-related cell loss via endoreplication-mediated cell growth. Nat Commun 17.

Ferrandon, D., J. L. Imler, C. Hetru and J. A. Hoffmann, 2007 The Drosophila systemic immune response: sensing and signalling during bacterial and fungal infections. Nat Rev Immunol 7: 862–874.

Gene Ontology, C., 2026 The Gene Ontology knowledgebase in 2026. Nucleic Acids Res 54: D1779–D1792.

Hallgren, J., K. D. Tsirigos, M. D. Pedersen, J. J. Almagro Armenteros, P. Marcatili et al., 2022 DeepTMHMM predicts alpha and beta transmembrane proteins using deep neural networks. bioRxiv: 2022.2004.2008.487609.

Handke, B., I. Poernbacher, S. Goetze, C. H. Ahrens, U. Omasits et al., 2013 The hemolymph proteome of fed and starved Drosophila larvae. PLoS One 8: e67208.

Hu, Y., A. Comjean, H. Attrill, G. Antonazzo, J. Thurmond et al., 2023 PANGEA: a new gene set enrichment tool for Drosophila and common research organisms. Nucleic Acids Res 51: W419–W426.

Hu, Y., A. Comjean, C. Gao, A. Veal, S. Yamamoto et al., 2026 DIOPT: the DRSC Integrative Ortholog Prediction Tool, 2026 update. Genetics.

Hu, Y., I. Flockhart, A. Vinayagam, C. Bergwitz, B. Berger et al., 2011 An integrative approach to ortholog prediction for disease-focused and other functional studies. BMC Bioinformatics 12: 357.

Kall, L., A. Krogh and E. L. Sonnhammer, 2007 Advantages of combined transmembrane topology and signal peptide prediction--the Phobius web server. Nucleic Acids Res 35: W429–432.

Kalra, H., R. J. Simpson, H. Ji, E. Aikawa, P. Altevogt et al., 2012 Vesiclepedia: a compendium for extracellular vesicles with continuous community annotation. PLoS Biol 10: e1001450.

Kanca, O., J. Zirin, Y. Hu, B. Tepe, D. Dutta et al., 2022 An expanded toolkit for Drosophila gene tagging using synthesized homology donor constructs for CRISPR-mediated homologous recombination. Elife 11.

Karlsson, M., C. Zhang, L. Mear, W. Zhong, A. Digre et al., 2021 A single-cell type transcriptomics map of human tissues. Sci Adv 7.

Kim, A. R., A. Comjean, A. Veal, J. Rodiger, M. Han et al., 2026 FlyPredictome: A structural atlas of predicted protein-protein interactions in Drosophila. bioRxiv.

Klee, E. W., 2008 The zebrafish secretome. Zebrafish 5: 131–138.

Krause, S. A., G. Overend, J. A. T. Dow and D. P. Leader, 2022 FlyAtlas 2 in 2022: enhancements to the Drosophila melanogaster expression atlas. Nucleic Acids Res 50: D1010–D1015.

Krogh, A., B. Larsson, G. von Heijne and E. L. Sonnhammer, 2001 Predicting transmembrane protein topology with a hidden Markov model: application to complete genomes. J Mol Biol 305: 567–580.

Kurusu, M., A. Cording, M. Taniguchi, K. Menon, E. Suzuki et al., 2008 A screen of cell-surface molecules identifies leucine-rich repeat proteins as key mediators of synaptic target selection. Neuron 59: 972–985.

Lane, E. A., A. Petsakou, Y. Liu, W. Chen, M. Qadiri et al., 2026 Cholinergic signaling modulates intestinal pathophysiology in a Drosophila model of cystic fibrosis. PLoS Genet 22: e1012048.

Lee, J. R., S. Urban, C. F. Garvey and M. Freeman, 2001 Regulated intracellular ligand transport and proteolysis control EGF signal activation in Drosophila. Cell 107: 161–171.

Li, H., J. Janssens, M. De Waegeneer, S. S. Kolluru, K. Davie et al., 2022 Fly Cell Atlas: A single-nucleus transcriptomic atlas of the adult fruit fly. Science 375: eabk2432.

Li, J., K. Huang, I. Dibra, Y. Liu, N. Perrimon et al., 2025 Desaturase-dependent secretory functions of hepatocyte-like cells control systemic lipid metabolism during starvation in Drosophila. Nat Commun 16: 10409.

Liu, Y., E. Dantas, M. Ferrer, T. Miao, M. Qadiri et al., 2025 Hepatic gluconeogenesis and PDK3 upregulation drive cancer cachexia in flies and mice. Nat Metab 7: 823–841.

Liu, Y., J. S. S. Li, J. Rodiger, A. Comjean, H. Attrill et al., 2022 FlyPhoneDB: an integrated web-based resource for cell-cell communication prediction in Drosophila. Genetics 220.

Lu, T. C., M. Brbic, Y. J. Park, T. Jackson, J. Chen et al., 2023 Aging Fly Cell Atlas identifies exhaustive aging features at cellular resolution. Science 380: eadg0934.

Marianes, A., and A. C. Spradling, 2013 Physiological and stem cell compartmentalization within the Drosophila midgut. Elife 2: e00886.

Mohr, S. E., Y. Hu, K. Kim, B. E. Housden and N. Perrimon, 2014 Resources for functional genomics studies in Drosophila melanogaster. Genetics 197: 1–18.

Nielsen, H., 2025 Practical Applications of Language Models in Protein Sorting Prediction: SignalP 6.0, DeepLoc 2.1, and DeepLocPro 1.0. Methods Mol Biol 2941: 153–175.

Onesto, M. M., C. A. Short, S. K. Rempel, T. S. Catlett and T. M. Gomez, 2021 Growth Factors as Axon Guidance Molecules: Lessons From in vitro Studies. Front Neurosci 15: 678454.

Ozturk-Colak, A., S. J. Marygold, G. Antonazzo, H. Attrill, D. Goutte-Gattat et al., 2024 FlyBase: updates to the Drosophila genes and genomes database. Genetics 227.

Palade, G., 1975 Intracellular aspects of the process of protein synthesis. Science 189: 347–358.

Pandey, U. B., and C. D. Nichols, 2011 Human disease models in Drosophila melanogaster and the role of the fly in therapeutic drug discovery. Pharmacol Rev 63: 411–436.

Pathan, M., P. Fonseka, S. V. Chitti, T. Kang, R. Sanwlani et al., 2019 Vesiclepedia 2019: a compendium of RNA, proteins, lipids and metabolites in extracellular vesicles. Nucleic Acids Res 47: D516–D519.

Pei, J., L. N. Kinch and N. V. Grishin, 2018 FlyXCDB-A Resource for Drosophila Cell Surface and Secreted Proteins and Their Extracellular Domains. J Mol Biol 430: 3353–3411.

Perkins, L. A., L. Holderbaum, R. Tao, Y. Hu, R. Sopko et al., 2015 The Transgenic RNAi Project at Harvard Medical School: Resources and Validation. Genetics 201: 843–852.

Petsakou, A., Y. Liu, Y. Liu, A. Comjean, Y. Hu et al., 2023 Cholinergic neurons trigger epithelial Ca(2+) currents to heal the gut. Nature 623: 122–131.

Qadiri, M., Y. Liu, A. R. Kim, M. Han, E. Zhou et al., 2025 FlyPhoneDB2: A computational framework for analyzing cell-cell communication in Drosophila scRNA-seq data integrating AlphaFold-multimer predictions. Comput Struct Biotechnol J 27: 2814–2822.

Saavedra, P., P. A. Dumesic, Y. Hu, E. Filine, P. Jouandin et al., 2023 REPTOR and CREBRF encode key regulators of muscle energy metabolism. Nat Commun 14: 4943.

Teufel, F., J. J. Almagro Armenteros, A. R. Johansen, M. H. Gislason, S. I. Pihl et al., 2022 SignalP 6.0 predicts all five types of signal peptides using protein language models. Nat Biotechnol 40: 1023–1025.

Trachana, K., T. A. Larsson, S. Powell, W. H. Chen, T. Doerks et al., 2011 Orthology prediction methods: a quality assessment using curated protein families. Bioessays 33: 769–780.

Tsirigos, K. D., C. Peters, N. Shu, L. Kall and A. Elofsson, 2015 The TOPCONS web server for consensus prediction of membrane protein topology and signal peptides. Nucleic Acids Res 43: W401–407.

Uhlen, M., M. J. Karlsson, A. Hober, A. S. Svensson, J. Scheffel et al., 2019 The human secretome. Sci Signal 12.

UniProt, C., 2023 UniProt: the Universal Protein Knowledgebase in 2023. Nucleic Acids Res 51: D523–D531.

Urban, S., J. R. Lee and M. Freeman, 2001 Drosophila rhomboid-1 defines a family of putative intramembrane serine proteases. Cell 107: 173–182.

van Kempen, M., S. S. Kim, C. Tumescheit, M. Mirdita, J. Lee et al., 2024 Fast and accurate protein structure search with Foldseek. Nat Biotechnol 42: 243–246.

Vierstraete, E., P. Verleyen, A. De Loof and L. Schoofs, 2005 Differential proteomics for studying Drosophila immunity. Ann N Y Acad Sci 1040: 504–507.

Wang, R., C. Ren, T. Gao, H. Li, X. Bo et al., 2024 SEPDB: a database of secreted proteins. Database (Oxford) 2024.

Xu, J., Y. Liu, F. Yang, Y. Cao, W. Chen et al., 2024 Mechanistic characterization of a Drosophila model of paraneoplastic nephrotic syndrome. Nat Commun 15: 1241.

Yoshida, Y., S. Kashio and M. Miura, 2026 Proximal labeling of the Golgi secretome reveals fat body-derived humoral factors in Drosophila disc regeneration. J Biol Chem 302: 113198.

Zirin, J., Y. Hu, L. Liu, D. Yang-Zhou, R. Colbeth et al., 2020 Large-Scale Transgenic Drosophila Resource Collections for Loss- and Gain-of-Function Studies. Genetics 214: 755–767.

