## Supplemental Figures for "The *Drosophila* Secretome"

**Supplementary figure 1:** Assembly of *Drosophila* Secretome and The Conservation Analysis

(A) *Drosophila* secretome assembly from annotation/data, signal and subcellular localization prediction as well as exosome vesicle annotation. Green circle is for genes obtained from annotated resources or dataset, light blue is for genes from computational predictions while the dark blue is for genes from Vesiclepedia database (exosome vesicle). (B) Conservation considering protein length.

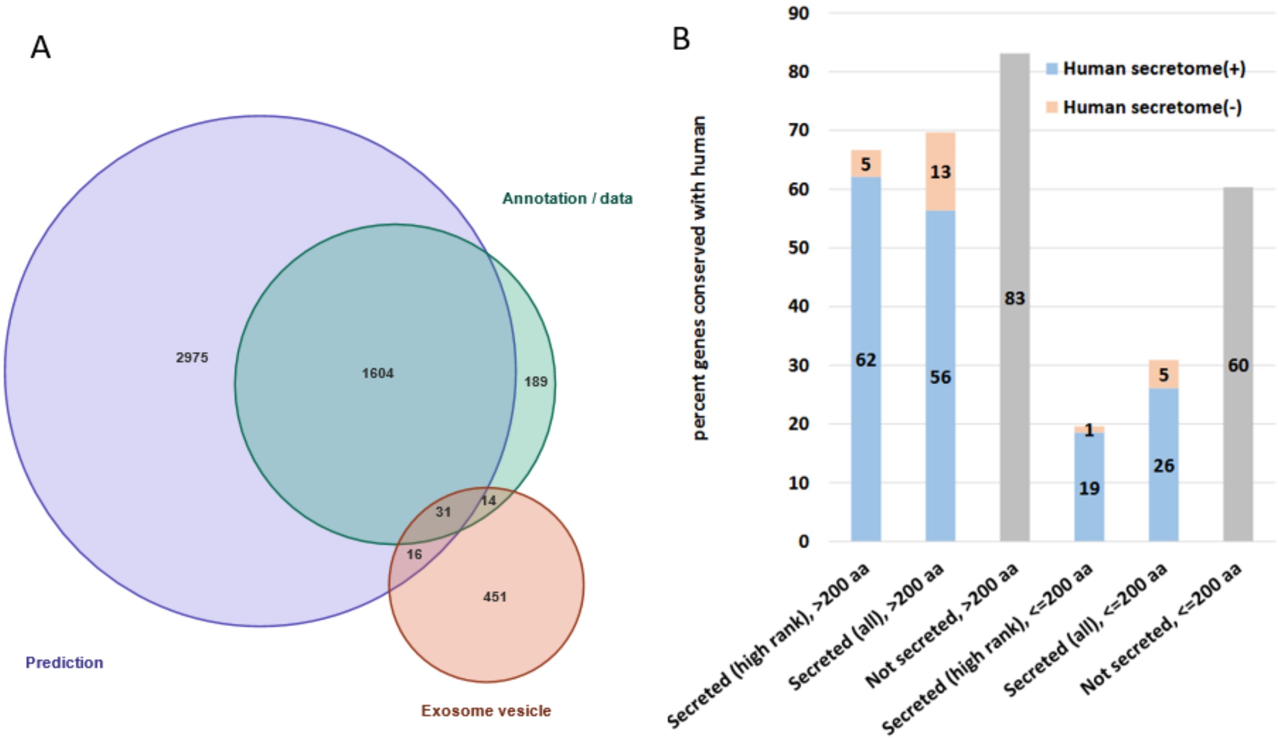

**Supplementary figure 2.** Tissue expression comparison between uncharacterized genes (CG genes) and the others (characterized genes) in *Drosophila* secretome.

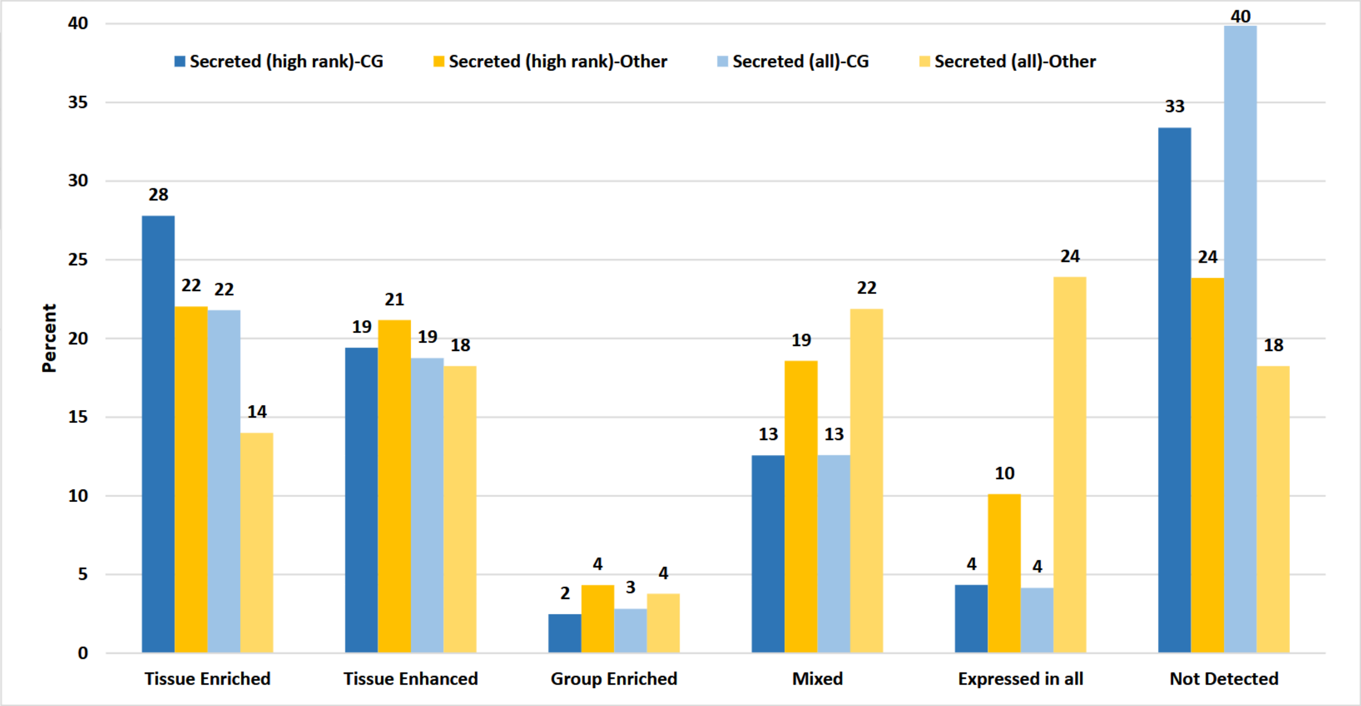

**Supplementary figure 3.** Tissue specific genes (tissue enriched and tissue enhanced) in various systems and their involvement in different biological processes. The color reflects the percentage of genes with a tissue-specific pattern vs all secreted genes for each major biological process listed.

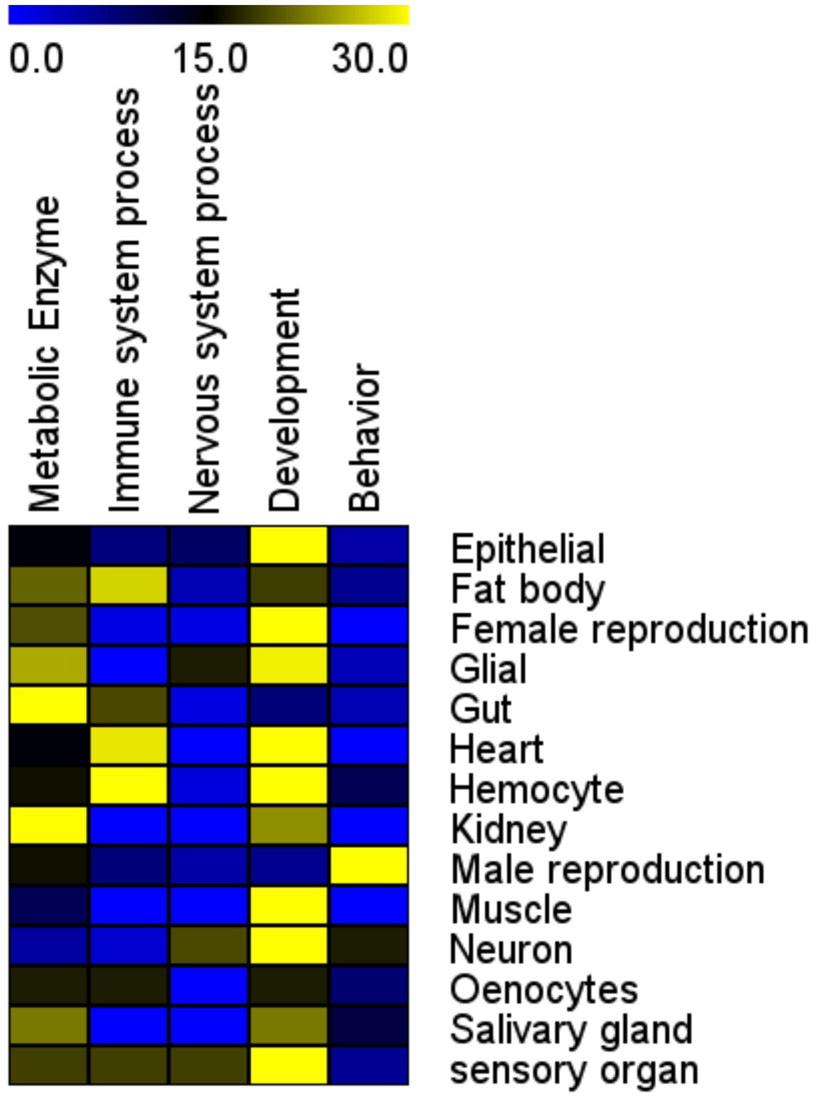

**Supplementary figure 4.** At *Drosophila* Secretome online portal, a menu bar with 13 tabs provides structured access to gene information, fly stock information, and other public resources, facilitating information mining and experimental design.

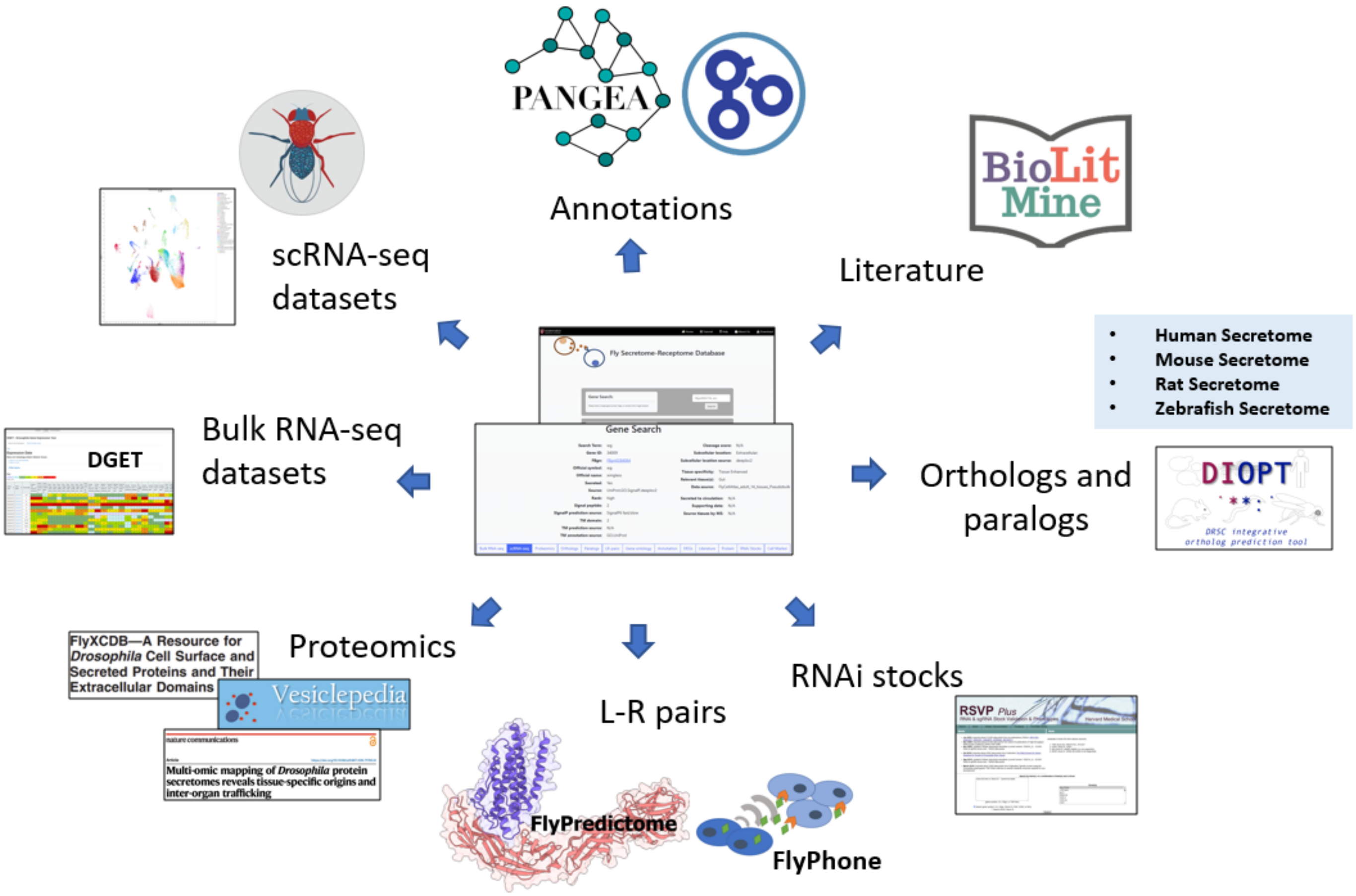
